# Membrane Insertion of *Mycobacterium tuberculosis* EsxA in Cultured Lung Epithelial Cells

**DOI:** 10.1101/2020.04.09.035238

**Authors:** Qi Zhang, Javier Aguilera, Salvador Vazquez Reyes, Jianjun Sun

## Abstract

EsxA has long been recognized as an important virulence factor of *Mycobacterium tuberculosis* (Mtb) that plays an essential role in Mtb cytosolic translocation presumably by penetrating phagosomal membranes with its acidic pH-dependent membrane permeabilizing activity (MPA). However, current data suggest that the observed cytolytic activity of EsxA at neutral pH is due to contamination of ASB-14, a detergent used in EsxA protein purification, and the role of EsxA MPA in Mtb cytosolic translocation is also questionable. Here, we have obtained evidence that it is ASB-14, not EsxA that causes cytolysis at neutral pH. Quantitative liquid chromatography and mass spectrometry showed that even after gel filtration, dialysis, or passing through detergent removal column, the remaining ASB-14 in the EsxA protein solution was still at a concentration enough to kill cultured lung epithelial cells. When treated with trypsin or proteinase K, the digested EsxA protein solution with ASB-14 was still cytotoxic. Interestingly, however, we have found that the exogenously added EsxA is endocytosed into lung epithelial cells and inserts into the host membranes within acidic subcellular compartments, which can be blocked by cytochalasin D and bafilomycin A. It is for the first time EsxA is found to insert into the host membranes within acidic subcellular compartments.

**Importance:** EsxA has long been recognized as an important virulence factor of *Mycobacterium tuberculosis* (Mtb) that plays an essential role in Mtb virulence. However, current data regarding to its role in Mtb virulence are controversial. Here, we have obtained evidence showing that the cytolytic activity of EsxA at neutral pH is due to contamination of ASB-14, a detergent used in EsxA preparation. Moreover, it is for the first time we have found that EsxA protein is endocytosed into lung epithelial cells and inserts into the host membranes within acidic subcellular compartments, implicating an important role of the acidic pH-dependent membrane permeabilizing activity of EsxA in Mtb virulence.

## Introduction

*Mycobacterium tuberculosis* (Mtb), the causative bacterial pathogen for tuberculosis disease, infects one-third of the world’s population and causes over one million deaths worldwide each year (1, 2). It is estimated that an airborne droplet carrying 1–3 Mtb bacilli is enough to cause systemic dissemination and infection (3, 4). When Mtb is inhaled into the lung, it is engulfed into alveolar macrophage. Instead of being destroyed in phagolysosomes like many other intracellular pathogens, Mtb develops effective strategies to evade host defense mechanisms. Recent studies have shown that Mtb is able to rupture the phagosomal membrane and translocate to the cytosol, where it replicates, eventually lyses the macrophage and spreads to the surrounding non-phagocytic cells, such as alveolar endothelial and epithelial cells (5–10). Thus, cytosolic translocation is regarded as one of the major mechanisms for Mtb virulence.

The ability of Mtb cytosolic translocation and dissemination is mainly dependent on ESX-1, the Type VII secretion system that is present in Mtb but deleted in the vaccine strain BCG (6, 8, 11–15). ESX-1 locus is comprised of 11 genes that encode the secretion apparatus and two secreted proteins: EsxA (6 kDa early secreted antigenic target, ESAT-6) and EsxB (10 kDa cell filtrate protein, CFP-10) (16–19). EsxA and EsxB are secreted in a co-dependent manner (20). Biochemical analysis has shown that EsxA and EsxB form a 1:1 heterodimeric complex (21–26). In our earlier studies, we have characterized the pH-dependent membrane interactions and conformational changes of EsxA and EsxB and found that at acidic pH (pH 5 and below) EsxA, but not EsxB, makes a significant conformational change and permeabilizes liposomal membranes (27). The membrane-permeabilizing activity (MPA) is unique to EsxA from Mtb, but not present in its ortholog from non-pathogenic *Mycobacterium smegmatis* (27). We have also showed that upon acidification the central Helix-Turn-Helix motif of EsxA inserts into the liposomal membrane and forms a membrane-spanning pore (28). Most recently, we have found glutamate 5 (Q5) is a key residue that regulates the MPA of EsxA (29). Mutations at Q5 either up- or down-regulated the MPA of EsxA *in vitro*, which accordingly either up- or down-regulated the mycobacterial cytosolic translocation and virulence in macrophages and zebra fishes (29). Thus, the data strongly support a model that within acidic subcellular compartments EsxA penetrates the host membranes and allows Mtb translocate to the cytosol for replication, eventually leading to macrophage lysis and bacterial dissemination to the surrounding cells and tissues (30). However, the direct evidence showing EsxA insertion into the host cell membrane is absent.

In addition to functioning within acidic subcellular compartments to facilitate mycobacterial cytosolic translocation, exogenously added EsxA protein has also been shown to cause cytolysis of red blood cells, macrophages, and type 1 and type 2 pneumocytes at neutral pH, suggesting that it may directly act on plasma membrane like a cytolytic pore-forming toxin (8, 21, 31–34). However, the data regarding to the ability of EsxA to cause cytolysis and cell death at neutral pH are controversial. In the studies assessing the effects of EsxA on immune responses, EsxA was shown to modulate the inflammatory responses of macrophages and T cells, but did not cause cytolysis or apoptosis (35–37). Refai et al reported that a zwitterionic detergent amidosulfobetaine-14 (ASB-14), which is commonly used in purification of recombinant EsxA in the protocol provided by BEI Resource, was involved in EsxA-mediated cytolysis (38). They found that the recombinant EsxA existed as a dimer or oligomer, which was not cytolytic, while the ASB-14-treated EsxA existed as a monomer with conformational changes and caused membrane-lysis and cell death (38). A recent study, however, showed that the recombinant EsxA protein didn’t lyse cell membrane and the lytic activity previously attributed to EsxA was due to the residual ASB-14 detergent in the preparation (39). They also found that Mm was still able to penetrate the phagosome and translocate to the cytosol in the presence of bafilomycin A, an intracellular acidification inhibitor, indicating that phagosome rupture doesn’t occur through the acidic pH-dependent MPA of EsxA (39). Most recently, Lienard et. al. employed a collection of Mm ESX-1 transposon mutants, including the mutants that disrupt EsxA secretion, to infect macrophages and showed that the transposon mutants without EsxA secretion was still able to permeabilize phagosomes (40). Thus, the data regarding to the role of EsxA MPA in mediating cytosolic translocation are conflicting. To solve the controversy, in the present study, we investigated the cytotoxic effects of ASB-14 and obtained evidence that it is ASB-14, not EsxA that causes cytolysis at neutral pH. More importantly, it is for the first time we found that EsxA was internalized into the cells through endocytosis, trafficked to acidic subcellular organelles and inserted into the host cell membrane.

## Materials and Methods

### Cell culture

The human type 1 alveolar epithelial-like cell line WI-26 was purchased from the American Type Culture Collection (ATCC CCL95.1), WI-26 cells are grown in MEM medium supplemented with 10% de-complemented FBS (Hyclone, Logan, UT).

### Growth of Mycobacterium marinum

Mm cells were grown in 7H9 liquid medium (Difco) supplemented with oleic acid-albumin-dextrose-catalase (OADC), 0.05% of glycerol and 0.02% Tween-80 at 30°C. At mid-log phase, the mycobacterial cells were harvested, washed, resuspended in PBS and then filtered through a 26-gauge needle syringe. Large clumps were further removed by centrifuge at 1000 × *g* for 2 min. The density of the single cell suspensions was measured by OD_600_.

### Protein expression and purification

EsxA was expressed and purified from *E. coli* BL-21(DE3) as described previously (27). Briefly, the inclusion body was isolated and then solubilized in 8 M urea. The proteins were refolded on a nickel column and eluted with an imidazole gradient. The eluted proteins were further purified by size exclusion chromatography using a Superdex 75 column that was pre-equilibrated with the buffer 20 mM TrisHCl, 100 mM NaCl, pH 7.0.

The EsxA mutants EsxA(S35C), EsxA(G10C) and EsxA(G88C) were generated by site-directed mutagenesis and purified as previously described (28). The EsxA(Q5K/S35C) was expressed and purified as a GST-fusion protein in BL-21 (DE3), followed by cleavage off the GST tag as previously described (27).

### Preparation of ASB-14-treated EsxAprotein

the purified EsxA protein was incubated with 0.5% ASB-14 overnight on ice. Then the samples were either applied to size exclusion chromatography using a Superdex 75 column or dialyzed in a Spectra/Por3 dialysis bag with 3 kDa MWCO in 20 mM Tris-Cl, 100 mM NaCl, pH 7.0.

### Fluorescence labeling

The EsxA proteins with Cys mutations were reduced with 20 mM dithiothreitol (DTT) on ice for 20 min. DTT was removed by passing through a Sephadex G-50 column that was pre-equilibrated in 20 mM Tris-Cl, 100 mM NaCl, pH 7.3. The proteins were concentrated to ~2 mg/ml using a 5-kDa-cutoff vivaspin concentrator (Vivascience). The proteins were incubated with 20-fold molar excess of either IANBD [*N*,*N*’-dimethyl-*N*-(iodoacetyl)-*N*’-(7-nitrobenz-2-oxa-1,3-diazol)ethylenediamine or Alexa Fluor 546 (AF546) maleimide (Molecular Probes] for 2 h at RT. The free dye was removed by passing through the G-50 Sephadex column. The fractions containing the NBD-labeled protein were pooled, concentrated and stored at −80 °C for future use. The labeling efficiency was calculated as [dye]/[protein]% by absorbance spectrophotometry (ε_478_ = 25,000 M^−1^ cm^−1^ for NBD, ε_554_ = 93,000 M^−1^ cm^−1^ for AF546, and ε_280_ = 17,739 M^−1^ cm^−1^ for EsxA). The measured labeling efficiency for all the mutants was ~ 100%.

### Detection of ASB-14 by LC-MS/MS

To establish the standard curve of ASB-14, ASB-14 standard (purchased from Sigma-Aldrich) was dissolved in MilliQ water at concentration of 0.01, 0.1, 1 and 10 mg/ml. The standard solution was run through a quantitative HPLC, followed by MS/MS to confirm the identity. To detect the ASB-14 contamination in the protein samples, the proteins were precipitated by adding 9 volumes of cold acetone and stored in −20 °C overnight. The samples were centrifuged at 15,000 × *g* for 30 min to precipitate the proteins, and the supernatant was transferred to a new tube, dried under nitrogen gas and dissolved in 0.1% AcF3. The samples were further desalted by a C18 desalting column and applied to a quantitative HPLC and MS/MS analysis.

### Time-lapse intensity measurements of NBD emission in live cells

WI-26 cells were detached by 0.25% trypsin and re-suspended into an universal buffer (10 mM HEPES, 10 mM sodium acetate, 10 mM MES, 150 mM NaCl, 2 mM CaCl2, 11 mM glucose, 50 mg/L bovine serum albumin, pH 7.3. The cells at OD_600_=0.05 (final concentration) were incubated with 100 μg NBD-labeled EsxA on ice for 1 h in a 2 ml total volume. The cells were then washed twice with cold universal buffer and transferred to a cuvette with a stirring bar in the cuvette holder of an ISS K2 fluorometer (ISS, IL). The temperature of the sample chamber was controlled by a circulating water bath. NBD was excited at 488 nm, and emission was recorded at 544 nm. In addition, a long-pass 510-nm filter was placed before the photomultiplier tube to reduce background scatter of the excitation beam. The total time to monitor NBD fluorescence change was 1 h. Relative Fluorescence change was calculated as the fluorescence intensity at the final time point subtracts the fluorescence intensity at 1,000 second when the temperature setting was changed from 4 °C to 37 °C. For the inhibitor experiments, the cells were incubated at 37 °C with 50 μM cytochalasin D or 1 μM bafilomycin A1 for 35 min before the NBD-labeled EsxA (S35C) protein was added to the cells.

### Confocal fluorescence microscopy

WI-26 cells were plated in a Lab-Tek chamber slide (Nalgen Nunc International, IL) at 3 × 10^4^ cells per well and incubated overnight. The slides were then incubated on ice, and a 1:1 mixture of the NBD-labeled EsxA and the AlexaFluro 546-labeled EsxA (Molecular Probes, OR) was added. After incubation for 1 h on ice, the slides were transferred to a humidified CO2 incubator at 37 °C. After 60 min, the cell samples were fixed for imaging and visualized under a Zeiss confocal microscope. The intensity of NBD was calibrated by the intensity of AF546.

## Results

### It is ASB-14, but not EsxA that causes cytolysis

To solve the controversy whether EsxA protein or ASB-14 causes cytolysis, we tested the potential cytotoxic effect of the purified EsxA protein in the presence or absence of ASB-14. The C-terminally His-tagged EsxA protein was overexpressed in the inclusion body in *E. coli*. The inclusion body was purified and solubilized by 8 M urea and then applied to a Ni-column, in which EsxA protein was refolded by slowly removing urea and eluted as a soluble protein with an imidazole gradient. No detergent is used during the purification process. We found that addition of the EsxA protein up to 40 μg/ml into the WI-26 cell culture caused little cytotoxicity (**Figure 1A**). Next, we mixed the EsxA protein with 0.5% ASB-14 overnight and then applied the protein-detergent mixture to gel filtration chromatography using a Superdex 75 column or to dialysis using 3 kDa cut-off filter membrane. We found that the resultant EsxA protein after gel filtration or dialysis caused significant cell death even at low protein concentrations (5-10 μg/ml) (**Figure 1A**).

**Fig 1.**
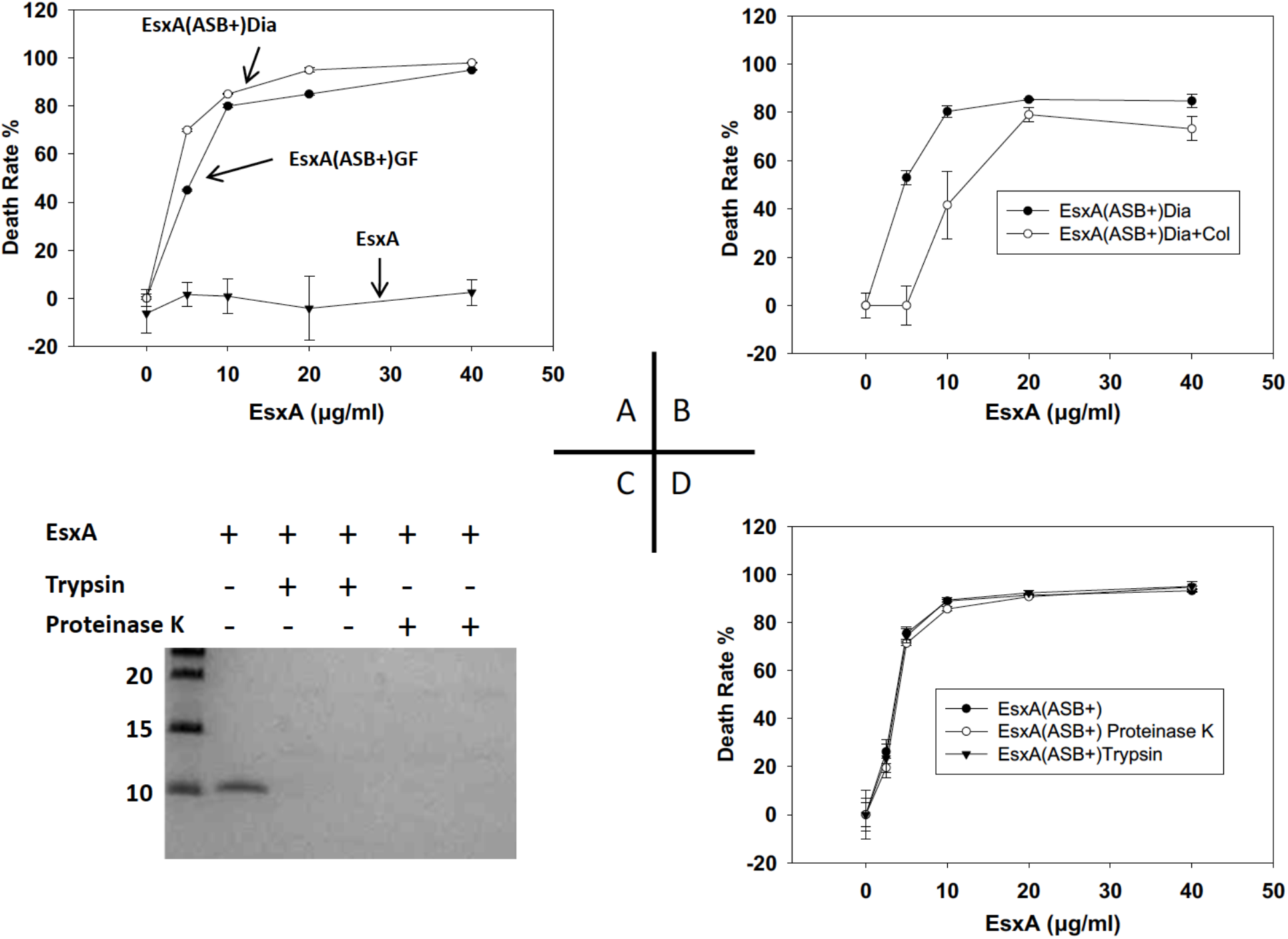
EsxA did not cause cytotoxicity in WI-26 cells. **A.** The purified EsxA protein was incubated with 0.5% ASB-14 for 4 h and then was either applied to gel filtration or dialysis to remove ASB-14. The resultant proteins were designated as EsxA(ASB+)GF or EsxA(ASB+)Dia. EsxA, EsxA(ASB+)GF and EsxA(ASB+)Dia were incubated with WI-26 cells at the indicated concentrations for 72 h. The cells that were incubated with medium alone (no protein) were set as negative controls. The cytotoxicity was measured by MTS assay and the death rate (%) was calculated as: (OD_492_ of control cells – OD_492_ of test cells)/(OD_492_ of test cells) X 100. The data were calculated from at least three independent experiments. **B**. The EsxA(ASB+)Dia sample was further passed through a HiPPR detergent removal column and the flow-through sample was collected and termed EsxA(ASB+)Dia+Col. EsxA(ASB+)Dia and EsxA(ASB+)Dia+Col were incubated with WI-26 for 72 h at the indicated concentrations. The cytotoxicity was measured as described above. **C**. The ASB-14-treated EsxA protein, EsxA(ASB+), was incubated with either 0.025% trypsin or 0.05% proteinase K for 30 min. The samples were applied to SDS-PAGE and stained by Coomarssie blue to confirm the digestion of EsxA protein after treatment of trypsin and proteinase K. **D.** Then the samples were incubated with WI-26 for 72 h. The cytotoxicity was measured as described above.

We suspected that ASB-14 was not completed removed by gel filtration or dialysis. Therefore, we applied the dialyzed EsxA protein sample to a HiPPR detergent removal column. Interestingly, while the sample treated with the HiPPR detergent removal column still caused cell death but had a significantly higher LD50 (~ 13 μg/ml), compared to the sample without passing through the HiPPR column (LD50 ~ 5 μg/ml) (**Figure 1B**). This indicates that the ASB-14 left in the EsxA protein solution may be responsible for cell death.

To further test our hypothesis, the EsxA protein solution containing 0.5% of ABS-14 was treated with trypsin or proteinase K. While SDS-PAGE showed that the EsxA protein was completely digested (**Figure 1C**), the digested protein solutions were still able to cause significant cell death (**Figure 1D**). Trypsin or proteinase K alone at the applied concentration did not cause any cell death (data not shown). These data strongly suggest that it is ASB-14, but not EsxA that caused cell death, and ASB-14 can’t be efficiently removed by gel filtration, dialysis and even the detergent removal column.

### The residual contamination of ASB-14 in EsxA protein preparation is enough to cause cell lysis

To further confirm that ASB-14 is cytotoxic to the cells, we tested the dose-dependent cytotoxicity of ASB-14. As low as 5 μg/ml, ASB-14 caused over 60% cell death, and at 50 μg/ml ASB-14 caused 100% cell death (**Figure 2A**). Now the question remains: after gel filtration, dialysis or passing through the HiPPR column, was the ASB-14 left in the protein solution enough to kill cells? We went ahead to determine the amount of ASB-14 in the EsxA protein solutions by quantitative HPLC. We first established a standard curve using the pure ASB-14 at various concentrations (**Figure 2B**) and then measured the abundance of ASB-14 in the EsxA protein solutions (**Figure 2C**). The identity of ASB-14 in protein solutions was confirmed by mass spectrometry. Based on the standard curve, the concentrations of ASB-14 in the protein solutions were calculated (**Table 1**). We further calculated the actual concentrations of ASB-14 that were applied into the tissue culture when the EsxA protein was used at 20 μg/ml (**Table 1**), a condition that killed at least 80% of cells (**Figure 1A, B and D**). The actual concentrations of ASB-14 applied in the cell culture ranged from 12-32 μg/ml (**Table 1**), which were enough to cause significant cytotoxicity as shown in **Figure 2A**. Together, the data confirmed that it was ASB-14, but not EsxA that caused cell death.

It is worth of mentioning that we tested ASB-14 compound from three different vendors (Sigma Aldrich, Santa Cruz Biotechnology, and EMD Millipore), and all brands of ASB-14 showed similar cytotoxic effects. We also purified the EsxA protein by following exactly the protocol from BEI Resource, and we found the EsxA protein solution was cytotoxic.

**Fig 2.**
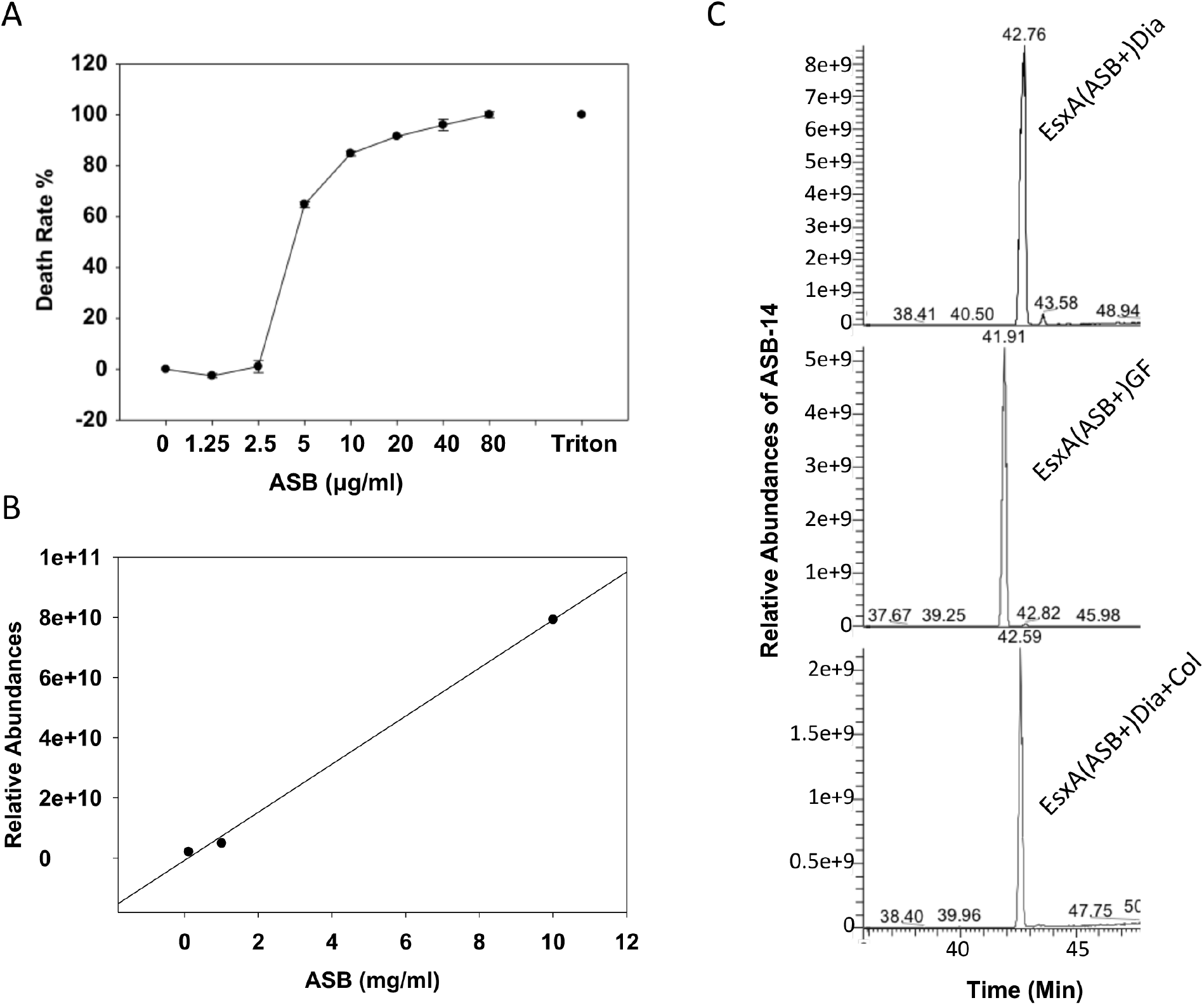
After gel filtration, dialysis, or passing through detergent removal column, the EsxA protein solution still has residual ASB-14 that is enough to kill cells. **A**. WI-26 cells were incubated with the standard solutions of ASB-14 at the indicated concentrations for 72 h. 0.1% Triton X-100 was used as a positive control. The death rate was measured and calculated as described above. **B**. The standard solutions of ASB-14 at various concentrations (0.001, 0.01, 0.1, 1, 10 mg/ml) were applied to the quantitative HPLC to establish the standard curve. **C**. The protein samples EsxA(ASB+)Dia, EsxA(ASB+)GF and EsxA(ASB+)Dia+Col were treated with 90% acetone at cold to precipitate proteins, and then the remaining acetone solutions were dried under nitrogen and the samples were applied to quantitative HPLC to quantify the abundance of ASB-14.

**Table 1:** Concentrations of ASB-14 in EsxA protein solutions measured by quantitative liquid chromatography

| Samples | ASB-14 concentration | Final concentration of ASB-14* |
| --- | --- | --- |
| EsxA(ASB+)GF | 1.46 mg/ml (3.36 mM) | 22.5 ug/ml (51.76 μM) |
| EsxA(ASB+)Dia | 1.93 mg/ml (4.44 mM) | 32.2 ug/ml (74.08 μM) |
| EsxA(ASB+)Dia+Col | 0.60 mg/ml (1.38 mM) | 12 ug/ml (27.6 μM) |

\*: The final concentrations of ASB-14 in the cultured cells, when EsxA is 20 μg/ml (2 $\mu$ M).

### EsxA protein was internalized into the host cell through endocytosis and inserted into the membrane under an acidic condition

Since EsxA is not cytolytic, we are intrigued to investigate whether EsxA can traffic into the cells and/or insert into the host membranes. NBD, an environmental sensitive dye, has little fluorescence in aqueous solution, but emits strong fluorescence when it inserts into lipid membranes. Thus, NBD has been used as a fluorescence marker for protein membrane insertion (41–43). Earlier, we have obtained evidence that the central Helix-Turn-Helix motif of EsxA inserts into the liposomal membrane at low pH (28). For instance, when NBD was labeled at S35C, a position that is embedded in the membrane, the NBD-labeled EsxA(S35C) showed a strong fluorescence upon acidification. However, when NBD was labeled at G10C, a position in the N-terminal arm that does not insert into the membrane, the NBD-labeled EsxA(G10C) did not emit fluorescence (28). Here, we used the NBD-labeled EsxA(S35S) and EsxA(G10C) to test the membrane insertion of EsxA within the host cells (**Figure 3**). The NBD-labeled proteins were first incubated with the cells at 4 °C, which allows the proteins bind to the cell surface without endocytosis. Then the temperature was shifted to 37 °C, which allows endocytosis to occur. The real-time fluorescence emission of the NBD-labeled proteins was monitored using a fluorometer. We found that upon temperature shifted from 4 °C to 37 °C, the fluorescence emission of EsxA(S35C)-NBD exhibited a significant increase over time, while EsxA(G10C)-NBD emitted little fluorescence, which is consistent to the result obtained in liposomes (28). Our recent study has identified Q5 as a key residue that regulates EsxA MPA (29). The mutation Q5K significantly reduced the MPA of EsxA and hence attenuated mycobacterial cytosolic translocation and virulence. Consistent to the result, we found that EsxA(Q5K/S35C)-NBD had a significantly lower fluorescence emission inside the cells (**Figure 3**). As expected, addition of cytochalasin D, which inhibits endocytosis by blocking actin polymerization, significantly reduced the fluorescence emission of EsxA(S35C)-NBD. Bafilomycin A, an inhibitor of endocytic acidification, exhibited a similar inhibitory effect. Together, the data suggest that EsxA is internalized into the host cell through endocytosis and inserts into the endosomal membranes in an acidic condition.

**Fig 3.**
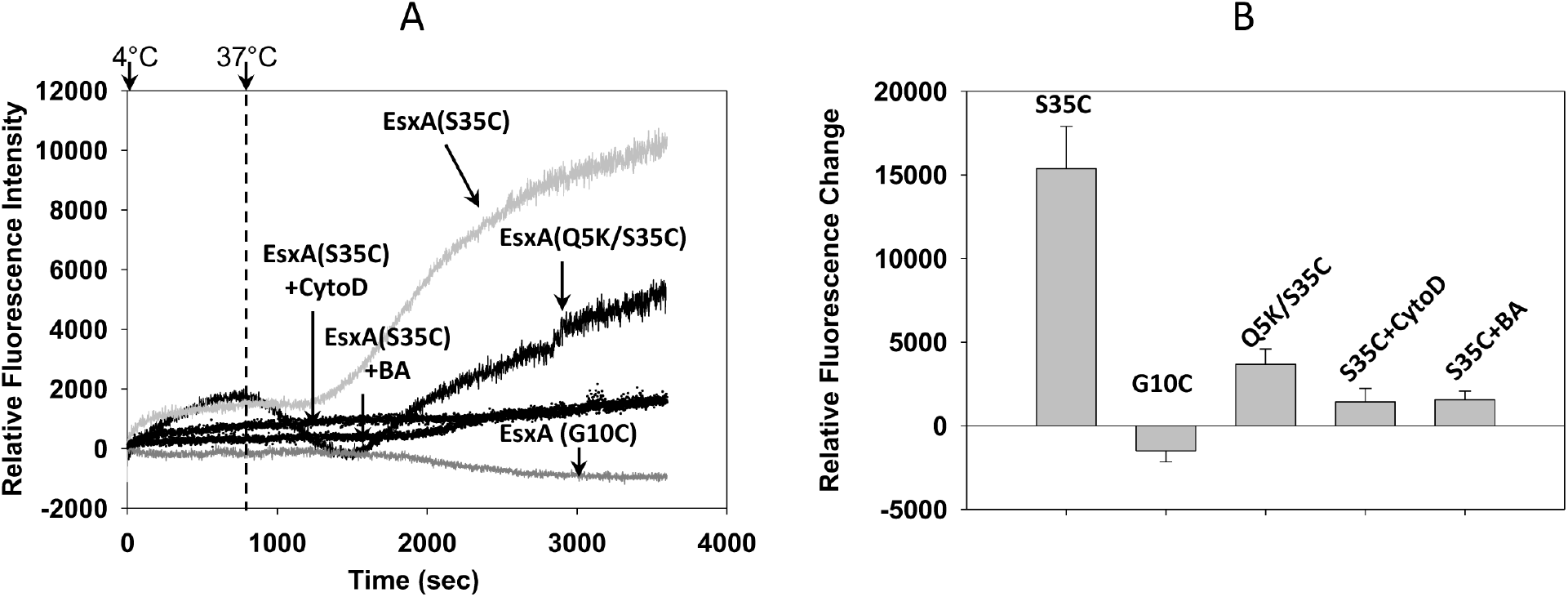
EsxA was internalized into the WI-26 cells through endocytosis and inserted into the endosomal membrane. The suspended WI-26 cells were incubated on ice with the NBD-labeled EsxA(S35C), EsxA(Q5K/S35C) or EsxA(G10C), respectively. After 30 min of incubation, the cells were washed and suspended in cold PBS and transferred to a cuvette with magnetic stirring in the cuvette chamber of a fluorometer. The cuvette chamber was pre-chilled at 4 °C by connecting to circulating water bath. At time zero, the temperature of the water bath was set to 37 °C, and the fluorescence intensity of NBD (Ex at 488 nm, Em at 545 nm) was recorded with time. To inhibit endocytosis, WI-26 cells were pre-treated with 50 μ M cytochalasin D (CytoD) or 1 μ M bafilomycin A1 (BA) for 35 min at 37 °C. The cells were then incubated with the NBD-labeled EsxA protein as described above. The representative NBD emission curves with time are shown in **A**. The relative increase of NBD fluorescence intensity at 1 h (3600 seconds) incubation was shown in **B**. The data were calculated from three independent experiments.

### Confocal fluorescence microscopy observed EsxA insertion into the host membrane within the acidic compartments

To directly visualize membrane insertion of EsxA inside the cells, we monitored the intracellular trafficking of EsxA(S35C)-NBD and EsxA(G10C)-NBD by confocal fluorescence microscopy (**Figure 4**). NBD emits green fluorescence when it inserts into the lipid membranes. To quantify the relative NBD fluorescence intensity of EsxA(S35C)-NBD and EsxA(G10C)-NBD, we used the AF546-labeled EsxA(G88C), which constitutively emits red fluorescence, as an internal reference of fluorescence intensity. Equal amount of EsxA(G88C)-AF546 was pre-mixed with EsxA(S35C)-NBD and EsxA(G10C)-NBD, respectively, followed by incubation with the cells. Thus, EsxA(G88C)-AF546 was co-trafficked with the NBD-labeled proteins, and its intensity is used to normalize the NBD fluorescence. Consistent to the result obtained in the fluorometer (**Figure 3**), the fluorescence intensity of EsxA(S35C)-NBD was significantly higher than that of EsxA(G10C)-NBD, which confirms the membrane insertion of EsxA within the host cells (**Figure 4A, B**).

**Fig 4.**
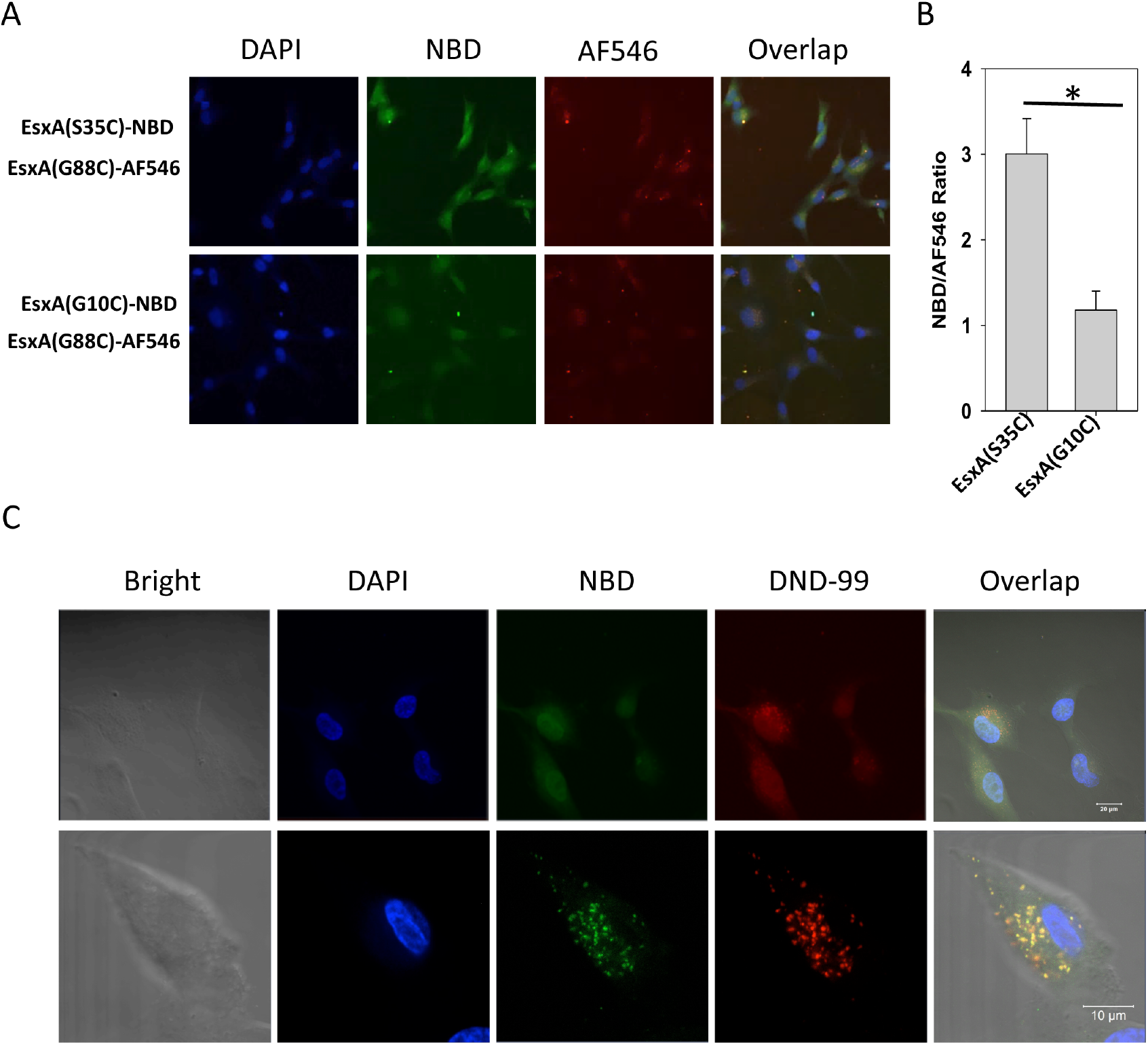
Fluorescence confocal microscopy detected membrane insertion of EsxA and its colocalization with the acidic lysotracker. **A**. The WI-26 cells were incubated on ice with either a 1:1 mixture of EsxA(S35C)-NBD and EsxA(G88C)-AF546 or a 1:1 mixture of EsxA(G10C)-NBD and EsxA(G88C)-AF546 for 1 h. The cells were washed and then transferred to a CO2 incubator at 37°C for 1 h. The cells were then fixed with 4% PFA and stained with DAPI. Finally, the cells were imaged under a fluorescence confocal microscope at three separate channels, blue (DAPI), green (NBD) and red (AF546). **B**. The intensities of NBD emission from EsxA(S35C) and EsxA(G10C) were measured and averaged from at least 6 random fields and the NBD intensity was normalized using the intensity of EsxA(G88C)-AF546 as an internal reference. The data was obtained from three independent experiments and were presented as mean ± S.E. (n = 3, *p < 0.05). **C.** WI-26 cells were incubated on ice with EsxA(S35C)-NBD and DND-99 for 1 h. The cells were washed and transferred to a CO2 incubator at 37°C for 1 h. The cells were fixed and stained with DAPI, followed by imaging under the confocal microscope using three channels: blue (DAPI), green (NBD) and red (DND-99).

To test if membrane insertion of EsxA occurs within the acidic subcellular organelles, we incubated the cells with EsxA(S35C)-NBD and stained the cells with LysoTracker DND-99, a red-fluorescent dye that is highly selective for acidic subcellular organelles (**Figure 4C**). At a higher magnification, both green fluorescence and red fluorescence exhibited as punctate spots and these spots were perfectly overlapped, suggesting that EsxA inserts into the membranes within acidic subcellular organelles (**Figure 4C**).

## Discussion

In our earlier biochemical characterization, we have found that EsxA requires acidic pH (≤ 5) to permeabilize liposomal membranes (27, 28). Recently, we have characterized the single-residue mutations at Q5 of EsxA (e.g. Q5K and Q5V) and demonstrated that the acidic pH-dependent MPA of EsxA is required for mycobacterial cytosolic translocation and virulence (29). Most recently, we have found that the N^α^-acetylation of EsxA at T2 is essential for the low pH-dependent EsxAB heterodimer dissociation, which is the prerequisite for EsxA membrane insertion. Single-residue mutations at T2 (e.g. T2A and T2R) disrupted N^α^-acetylation, resulting in attenuated cytosolic translocation and virulence (44). All these findings support that EsxA MPA functions within the acidic subcellular compartments and contributes to the virulence of Mtb through rupturing the host phagosome membranes. Interestingly, several studies have reported that the exogenously added recombinant EsxA acted directly on plasma membrane and caused cytolysis at neutral pH, suggesting that EsxA could function as a cytolytic pore-forming toxin (8, 21, 31–34). On the contrary, however, several other studies didn’t observe any cytolytic effect of the exogenously added EsxA on the cultured mammalian cells (35–37). In the present study, we further investigated this discrepancy and obtained evidence that ASB-14, the detergent used in EsxA preparation, is responsible for the observed cytolysis. ASB-14 is a zwitterionic detergent and useful for extraction of membrane proteins. ASB-14 has molecular weight 434.68 and critical micelle concentration at 8 mM. When 0.5% (11.5 mM) of ASB-14 was used in EsxA purification, even after gel filtration, dialysis and passage through the HiRPP detergent removal column, there was still significant amount of ASB-14 (1.38 – 4.44 mM) present in the protein solution (**Table 1**). By comparing the ASB-14 cytotoxicity titration curve, the residual ASB-14 was enough to cause significant cytolysis (**Figure 2**). Another evidence comes from the digestion experiment. Even though EsxA was completely digested by the proteases, the solution containing ASB-14 was still cytotoxic (**Figure 1C, D**). Therefore, this study has further clarified the controversy and demonstrated that EsxA doesn’t cause cytolysis on plasma membrane at neutral pH. This is consistent with our earlier studies that EsxA requires acidic pH to permeabilize the membrane (27).

Instead of causing cytolysis, we found that EsxA was internalized into the host cell through endocytosis and trafficked to the acidic subcellular organelles, where it inserts into the membranes. To the best of our knowledge, it is for the first time that EsxA acidic pH-dependent membrane insertion is observed inside the host cells, which is consistent to the results obtained in liposomes (27–29, 45).

Our previous study has found that deletion of the *esxB-esxA* operon in Mm significantly reduced the adherence of Mm to the pre-fixed macrophages, and complementation of *esxB-esxA* restored the adherence, indicating that EsxA and/or EsxB mediate adherence of mycobacteria to the host cells through binding to the receptors on the host cell surface (29). The results obtained in the present study further indicate the presence of receptor molecule(s) on the host cell surface that binds to EsxA and takes it into specific endocytic pathways. While the surface receptors for EsxA and/or EsxB have not been identified yet, several earlier studies have presented evidence that EsxA or the heterodimer interacted with various host proteins. Earlier, Renshaw et al. found that the fluorescently labeled EsxAB heterodimer bound to the surfaces of monocytes and macrophages, but not fibroblasts (25). Later, EsxA was shown to directly bind to TLR2, which inhibited TLR signaling in macrophages (37). In a separate study, however, EsxA was shown to bind to type 1 and type 2 pneumocytes and purified human laminin (33). Recently, in a yeast two-hybrid screening, EsxA was found to bind directly to β2-microglobulin (β2M) and enter ER where it sequestered β2M to downregulate class-I-mediated antigen presentation (46).

It is generally believed that EsxAB is secreted out of mycobacteria, but the results about its localization are still controversial. In current studies, EsxAB has been localized to mycobacterial cell wall, cell surface, capsule and bacteriological medium (13, 19, 33, 47, 48). Champion et al. have showed that some EsxA molecules remained bound to the surface of mycobacteria and the surface-associated EsxA caused more cytotoxicity than those in the supernatant (49, 50). Mycobacteria caused the ESX-1-mediated hemolysis of red blood cells in a contact-dependent manner (39). Phthiocerol dimycocerosates (PDIM), a lipid virulence factor on mycobacterial cell wall, has been shown to act in concert with EsxA to cause phagosomal rupture and host cell apoptosis (51). All of the evidence mentioned above support that EsxAB is located on mycobacterial cell wall, where it acts in concert with other components of the cell wall (e.g. PDIM), binds to the specific receptor(s) of host cell, and brings mycobacterial cell wall and host cell membrane to close proximity, facilitating mycobacterial phagocytosis and subsequent cytosolic translocation through EsxA-mediated phagosome rupture.

In future, more in-depth studies are needed to investigate the mechanism of EsxA-mediated pathogen-host interaction, such as identification of the receptors and intracellular trafficking pathways.

## Acknowledgement

The study is supported by the grants from NIGMS (SC1GM095475 to J. Sun), National Center for Research Resources (5G12RR008124) and National Institute on Minority Health and Health Disparities (G12MD007592).

## References

1. Dolin, P. J., Raviglione, M. C., and Kochi, A. (1994) Global tuberculosis incidence and mortality during 1990-2000. Bull. World Health Organ. 72, 213–220

2. WHO (2015) Global tuberculosis report 2015

3. Birkness, K. A., Deslauriers, M., Bartlett, J. H., White, E. H., King, C. H., and Quinn, F. D. (1999) An in vitro tissue culture bilayer model to examine early events in Mycobacterium tuberculosis infection. Infect Immun. 67, 653–658

4. Humphreys, I. R., Stewart, G. R., Turner, D. J., Patel, J., Karamanou, D., Snelgrove, R. J., and Young, D. B. (2006) A role for dendritic cells in the dissemination of mycobacterial infection. Microbes Infect. 8, 1339–1346

5. van der Wel, N., Hava, D., Houben, D., Fluitsma, D., van Zon, M., Pierson, J., Brenner, M., and Peters, P. J. (2007) M. tuberculosis and M. leprae translocate from the phagolysosome to the cytosol in myeloid cells. Cell. 129, 1287–1298

6. Houben, D., Demangel, C., van Ingen, J., Perez, J., Baldeón, L., Abdallah, A. M., Caleechurn, L., Bottai, D., van Zon, M., De Punder, K., van der Laan, T., Kant, A., Bossers-de Vries, R., Willemsen, P., Bitter, W., van Soolingen, D., Brosch, R., van der Wel, N., and Peters, P. J. (2012) ESX-1 Mediated Translocation to the Cytosol controls Virulence of Mycobacteria. Cell Microbiol. 14, 1287–1298

7. Dobos, K. M., Spotts, E. A., Quinn, F. D., and King, C. H. (2000) Necrosis of lung epithelial cells during infection with Mycobacterium tuberculosis is preceded by cell permeation. Infect Immun. 68, 6300–6310

8. Hsu, T., Hingley-Wilson, S. M., Chen, B., Chen, M., Dai, A. Z., Morin, P. M., Marks, C. B., Padiyar, J., Goulding, C., Gingery, M., Eisenberg, D., Russell, R. G., Derrick, S. C., Collins, F. M., Morris, S. L., King, C. H., and William R Jacobs, J. (2003) The primary mechanism of attenuation of bacillus Calmette–Guérin is a loss of secreted lytic function required for invasion of lung interstitial tissue. Proc Natl Acad Sci USA. 100, 12420–12425

9. Simeone, R., Bobard, A., Lippmann, J., Bitter, W., Majlessi, L., Brosch, R., and Enninga, J. (2012) Phagosomal rupture by Mycobacterium tuberculosis results in toxicity and host cell death. PLoS Pathog. 8, e1002507

10. Simeone, R., Sayes, F., Song, O., Gröschel, M. I., Brodin, P., Brosch, R., and Majlessi, L. (2015) Cytosolic access of Mycobacterium tuberculosis: critical impact of phagosomal acidification control and demonstration of occurrence in vivo. PLoS Pathog. 11, e1004650

11. Mahairas, G. G., Sabo, P. J., Hickey, M. J., Singh, D. C., and Stover, C. K. (1996) Molecular analysis of genetic differences between Mycobacterium bovis BCG and virulent M. bovis. J Bacteriol. 178, 1274–1282

12. Harboe, M., Oettinger, T., Wiker, H. G., Rosenkrands, I., and Andersen, P. (1996) Evidence for occurrence of the ESAT-6 protein in Mycobacterium tuberculosis and virulent Mycobacterium bovis and for its absence in Mycobacterium bovis BCG. Infect Immun. 64, 16–22

13. Pym, A. S., Brodin, P., Brosch, R., Huerre, M., and Cole, S. T. (2002) Loss of RD1 contributed to the attenuation of the live tuberculosis vaccines Mycobacterium bovis BCG and Mycobacterium microti. Mol Microbiol. 46, 709–717

14. Stanley, S. A., Raghavan, S., Hwang, W. W., and Cox, J. S. (2003) Acute infection and macrophage subversion by Mycobacterium tuberculosis require a specialized secretion system. Proc Natl Acad Sci USA. 100, 13001–13006

15. Guinn, K. M., Hickey, M. J., Mathur, S. K., Zakel, K. L., Grotzke, J. E., Lewinsohn, D. M., Smith, S., and Sherman, D. R. (2004) Individual RD1-region genes are required for export of ESAT-6/CFP-10 and for virulence of Mycobacterium tuberculosis. Mol Microbiol. 51, 359–370

16. Simeone, R., Bottai, D., and Brosch, R. (2009) ESX/type VII secretion systems and their role in host-pathogen interaction. Curr Opin Microbiol. 12, 4–10

17. Andersen, P., Andersen, A. B., Sørensen, A. L., and Nagai, S. (1995) Recall of long-lived immunity to Mycobacterium tuberculosis infection in mice. J Immunol. 154, 3359–3372

18. Sørensen, A. L., Nagai, S., Houen, G., Andersen, P., and Andersen, A. B. (1995) Purification and characterization of a low-molecular-mass T-cell antigen secreted by Mycobacterium tuberculosis. Infect Immun. 63, 1710–1717

19. Berthet, F. X., Rasmussen, P. B., Rosenkrands, I., Andersen, P., and Gicquel, B. (1998) A Mycobacterium tuberculosis operon encoding ESAT-6 and a novel low-molecular-mass culture filtrate protein (CFP-10). Microbiology (Reading, Engl). 144 (Pt 11), 3195–3203

20. Fortune, S. M., Jaeger, A., Sarracino, D. A., Chase, M. R., Sassetti, C. M., Sherman, D. R., Bloom, B. R., and Rubin, E. J. (2005) Mutually dependent secretion of proteins required for mycobacterial virulence. Proc Natl Acad Sci USA. 102, 10676–10681

21. de Jonge, M. I., Pehau-Arnaudet, G., Fretz, M. M., Romain, F., Bottai, D., Brodin, P., Honoré, N., Marchal, G., Jiskoot, W., England, P., Cole, S. T., and Brosch, R. (2007) ESAT-6 from Mycobacterium tuberculosis dissociates from its putative chaperone CFP-10 under acidic conditions and exhibits membrane-lysing activity. J Bacteriol. 189, 6028–6034

22. Lightbody, K. L., Ilghari, D., Waters, L. C., Carey, G., Bailey, M. A., Williamson, R. A., Renshaw, P. S., and Carr, M. D. (2008) Molecular features governing the stability and specificity of functional complex formation by Mycobacterium tuberculosis CFP-10/ESAT-6 family proteins. J Biol Chem. 283, 17681–17690

23. Meher, A. K., Bal, N. C., Chary, K. V. R., and Arora, A. (2006) Mycobacterium tuberculosis H37Rv ESAT-6-CFP-10 complex formation confers thermodynamic and biochemical stability. FEBS J. 273, 1445–1462

24. Renshaw, P. S., Panagiotidou, P., Whelan, A., Gordon, S. V., Hewinson, R. G., Williamson, R. A., and Carr, M. D. (2002) Conclusive evidence that the major T-cell antigens of the Mycobacterium tuberculosis complex ESAT-6 and CFP-10 form a tight, 1:1 complex and characterization of the structural properties of ESAT-6, CFP-10, and the ESAT-6*CFP-10 complex. Implications for pathogenesis and virulence. J Biol Chem. 277, 21598–21603

25. Renshaw, P. S., Lightbody, K. L., Veverka, V., Muskett, F. W., Kelly, G., Frenkiel, T. A., Gordon, S. V., Hewinson, R. G., Burke, B., Norman, J., Williamson, R. A., and Carr, M. D. (2005) Structure and function of the complex formed by the tuberculosis virulence factors CFP-10 and ESAT-6. EMBO J. 24, 2491–2498

26. Lightbody, K. L., Renshaw, P. S., Collins, M. L., Wright, R. L., Hunt, D. M., Gordon, S. V., Hewinson, R. G., Buxton, R. S., Williamson, R. A., and Carr, M. D. (2004) Characterisation of complex formation between members of the Mycobacterium tuberculosis complex CFP-10/ESAT-6 protein family: towards an understanding of the rules governing complex formation and thereby functional flexibility. FEMS Microbiol Lett. 238, 255–262

27. De Leon, J., Jiang, G., Ma, Y., Rubin, E., Fortune, S., and Sun, J. (2012) Mycobacterium tuberculosis ESAT-6 Exhibits a Unique Membrane-interacting Activity That Is Not Found in Its Ortholog from Non-pathogenic Mycobacterium smegmatis. J Biol Chem. 287, 44184–44191

28. Ma, Y., Keil, V., and Sun, J. (2015) Characterization of Mycobacterium tuberculosis EsxA membrane insertion: roles of N-and C-terminal flexible arms and central helix-turn-helix motif. J Biol Chem. 290, 7314–7322

29. Zhang, Q., Wang, D., Jiang, G., Liu, W., Deng, Q., Li, X., Qian, W., Ouellet, H., and Sun, J. (2016) EsxA membrane-permeabilizing activity plays a key role in mycobacterial cytosolic translocation and virulence: effects of single-residue mutations at glutamine 5. Sci Rep. 6, 32618

30. Peng, X., and Sun, J. (2016) Mechanism of ESAT-6 membrane interaction and its roles in pathogenesis of Mycobacterium tuberculosis. Toxicon. 116, 29–34

31. Smith, J., Manoranjan, J., Pan, M., Bohsali, A., Xu, J., Liu, J., McDonald, K. L., Szyk, A., LaRonde-LeBlanc, N., and Gao, L.-Y. (2008) Evidence for pore formation in host cell membranes by ESX-1-secreted ESAT-6 and its role in Mycobacterium marinum escape from the vacuole. Infect Immun. 76, 5478–5487

32. Derrick, S. C., and Morris, S. L. (2007) The ESAT6 protein of Mycobacterium tuberculosis induces apoptosis of macrophages by activating caspase expression. Cell Microbiol. 9, 1547–1555

33. Kinhikar, A. G., Verma, I., Chandra, D., Singh, K. K., Weldingh, K., Andersen, P., Hsu, T., Jacobs, W. R., Jr, and Laal, S. (2010) Potential role for ESAT6 in dissemination of M.â□ □tuberculosisvia human lung epithelial cells. Mol Microbiol. 75, 92–106

34. Francis, R. J., Butler, R. E., and Stewart, G. R. (2014) Mycobacterium tuberculosis ESAT-6 is a leukocidin causing Ca2γ influx, necrosis and neutrophil extracellular trap formation. Cell Death Dis. 5, e1474

35. Peng, H., Wang, X., Barnes, P. F., Tang, H., Townsend, J. C., and Samten, B. (2011) The Mycobacterium tuberculosis early secreted antigenic target of 6 kDa inhibits T cell interferon-γ production through the p38 mitogen-activated protein kinase pathway. J Biol Chem. 286, 24508–24518

36. Chatterjee, S., Dwivedi, V. P., Singh, Y., Siddiqui, I., Sharma, P., Van Kaer, L., Chattopadhyay, D., and Das, G. (2011) Early secreted antigen ESAT-6 of Mycobacterium tuberculosis promotes protective T helper 17 cell responses in a toll-like receptor-2-dependent manner. PLoS Pathog. 7, e1002378

37. Pathak, S. K., Basu, S., Basu, K. K., Banerjee, A., Pathak, S., Bhattacharyya, A., Kaisho, T., Kundu, M., and Basu, J. (2007) Direct extracellular interaction between the early secreted antigen ESAT-6 of Mycobacterium tuberculosis and TLR2 inhibits TLR signaling in macrophages. Nat Immunol. 8, 610–618

38. Refai, A., Haoues, M., Othman, H., Barbouche, M. R., Moua, P., Bondon, A., Mouret, L., Srairi-Abid, N., and Essafi, M. (2015) Two distinct conformational states of Mycobacterium tuberculosis virulent factor early secreted antigenic target 6 kDa are behind the discrepancy around its biological functions. FEBS J. 10.1111/febs.13408

39. Conrad, W. H., Osman, M. M., Shanahan, J. K., Chu, F., Takaki, K. K., Cameron, J., Hopkinson-Woolley, D., Brosch, R., and Ramakrishnan, L. (2017) Mycobacterial ESX-1 secretion system mediates host cell lysis through bacterium contact-dependent gross membrane disruptions. Proc Natl Acad Sci USA. 114, 1371–1376

40. Lienard, J., Nobs, E., Lovins, V., Movert, E., Valfridsson, C., and Carlsson, F. (2019) The Mycobacterium marinum ESX-1 system mediates phagosomal permeabilization and type I interferon production via separable mechanisms. Proc Natl Acad Sci USA. 17, 201911646

41. Sun, J., Vernier, G., Wigelsworth, D. J., and Collier, R. J. (2007) Insertion of anthrax protective antigen into liposomal membranes: effects of a receptor. J Biol Chem. 282, 1059–1065

42. Sun, J., Lang, A. E., Aktories, K., and Collier, R. J. (2008) Phenylalanine-427 of anthrax protective antigen functions in both pore formation and protein translocation. Proc Natl Acad Sci USA. 105, 4346–4351

43. Jacquez, P., Avila, G., Boone, K., Altiyev, A., Puschhof, J., Sauter, R., Arigi, E., Ruiz, B., Peng, X., Almeida, I., Sherman, M., Xiao, C., and Sun, J. (2015) The Disulfide Bond Cys255-Cys279 in the Immunoglobulin-Like Domain of Anthrax Toxin Receptor 2 Is Required for Membrane Insertion of Anthrax Protective Antigen Pore. PLoS ONE. 10, e0130832

44. Nα-acetylation of the virulence factor EsxA is required for mycobacterial cytosolic translocation and virulence. (2020) Nα-acetylation of the virulence factor EsxA is required for mycobacterial cytosolic translocation and virulence. 10.1074/jbc.RA119.012497

45. Peng, X., Jiang, G., Liu, W., Zhang, Q., Qian, W., and Sun, J. (2016) Characterization of differential pore-forming activities of ESAT-6 proteins from Mycobacterium tuberculosis and Mycobacterium smegmatis. FEBS Lett. 590, 509–519

46. Sreejit, G., Ahmed, A., Parveen, N., Jha, V., Valluri, V. L., Ghosh, S., and Mukhopadhyay, S. (2014) The ESAT-6 Protein of Mycobacterium tuberculosis Interacts with Beta-2-Microglobulin (β2M) Affecting Antigen Presentation Function of Macrophage. PLoS Pathog. 10, e1004446

47. Majlessi, L., Brodin, P., Brosch, R., Rojas, M.-J., Khun, H., Huerre, M., Cole, S. T., and Leclerc, C. (2005) Influence of ESAT-6 secretion system 1 (RD1) of Mycobacterium tuberculosis on the interaction between mycobacteria and the host immune system. J Immunol. 174, 3570–3579

48. Sani, M., Houben, E. N. G., Geurtsen, J., Pierson, J., De Punder, K., van Zon, M., Wever, B., Piersma, S. R., Jiménez, C. R., Daffé, M., Appelmelk, B. J., Bitter, W., van der Wel, N., and Peters, P. J. (2010) Direct visualization by cryo-EM of the mycobacterial capsular layer: a labile structure containing ESX-1-secreted proteins. PLoS Pathog. 6, e1000794

49. Kennedy, G. M., Hooley, G. C., Champion, M. M., Mba Medie, F., and Champion, P. A. D. (2014) A Novel ESX-1 Locus Reveals that Surface-Associated ESX-1 Substrates Mediate Virulence in Mycobacterium marinum. J Bacteriol. 196, 1877–1888

50. Medie, F. M., Champion, M. M., Williams, E. A., and DiGiuseppe Champion, P. A. (2014) Homeostasis of N-α terminal acetylation of EsxA correlates with virulence in Mycobacterium marinum. Infect Immun. 10.1128/IAI.02153-14

51. Augenstreich, J., Arbues, A., Simeone, R., Haanappel, E., Wegener, A., Sayes, F., Chevalier, F. L., Chalut, C., Malaga, W., Guilhot, C., Brosch, R., and Astarie-Dequeker, C. (2017) ESX-1 and phthiocerol dimycocerosates of Mycobacterium tuberculosis act in concert to cause phagosomal rupture and host cell apoptosis. Cell Microbiol. 19, e12726

